# Proprioception-based movement goals support imitation and are disrupted in apraxia

**DOI:** 10.1101/2021.03.17.435845

**Authors:** Mitchell W. Isaacs, Laurel J. Buxbaum, Aaron L. Wong

## Abstract

The ability to imitate observed actions serves as an efficient method for learning novel movements and is specifically impaired (without concomitant gross motor impairments) in the neurological disorder of limb apraxia, a disorder common after left hemisphere stroke. Research with apraxic patients has advanced our understanding of how people imitate. However, the role of proprioception in imitation has been rarely assessed directly. Prior work has proposed that proprioceptively sensed body position is transformed into a visual format, supporting the attainment of a desired imitation goal represented visually (i.e., how the movement should look when performed). In contrast, we hypothesized a more direct role for proprioception: we suggest that movement goals are also represented proprioceptively (i.e., how a desired movement should feel when performed), and the ability to represent or access such proprioceptive goals is deficient in apraxia. Using a novel imitation task in which a robot cued meaningless trajectories proprioceptively or visually, we probed the role of each sensory modality. We found that patients with left hemisphere stroke were disproportionately worse than controls at imitating when cued proprioceptively versus visually. This proprioceptive versus visual disparity was associated with apraxia severity as assessed by a traditional imitation task, but could not be explained by general proprioceptive impairment or speed-accuracy trade-offs. These data suggest that successful imitation depends in part on the ability to represent movement goals in terms of how those movements should feel, and that deficits in this ability contribute to imitation impairments in patients with apraxia.

## 1 INTRODUCTION

Imitation is a valuable skill for rapidly learning novel actions. By directly copying an actor’s movements with one’s own body, imitation allows people to skip much of the trial-and-error process involved in learning to produce sophisticated behaviors such as making a layup in basketball. Surprisingly, despite over a century of study, it remains unclear exactly how imitation is accomplished. When we think of imitation, we typically assume that imitation occurs after visually observing others move; that is, we see what a movement is supposed to look like, then attempt to reproduce it using our own body. This has led to a common (albeit largely implicit) assumption that desired imitation goals are represented visually; that is, we plan a movement that will reproduce how an observed action looks. Specifying the desired action to be performed constitutes a movement goal (Passingham & Wise, 2012; Wong et al., 2015), which in this case is assumed to be represented in a visual format. In this context, the role of proprioceptive feedback has largely been speculated to simply support this process by contributing to the formation of an estimate of the current state of the body, e.g. by mapping kinesthetic input into an estimate of how the body appears visually (Meltzoff & Moore, 1977; Mitchell, 2002; Wolpert, 1997). This estimate of the current body state can then be compared against the desired movement goal to ensure movements are performed accurately (Diedrichsen et al., 2010; Zimmermann et al., 2012). Here, we question whether this is indeed the only role of proprioception in imitation.

The notion that imitation goals are specified visually has been supported by studies of patients with limb apraxia. Often arising from left-hemisphere strokes, apraxia frequently results in imitation impairments (for both static and dynamic movements) that may be observed even on the ipsilesional side in the absence of gross underlying motor deficits (Bartolo et al., 2008; Buxbaum, Johnson-Frey, et al., 2005; Buxbaum et al., 2014; Liepmann, 1905; Goldenberg, 2009, 2014; Goldenberg & Hagmann, 1997; Hermsdörfer et al., 1996; Rothi et al., 1991; Tarhan et al., 2015). In contrast, these patients tend to have relatively unimpaired reaching (e.g., aiming to a target in space) and grasping (Buxbaum et al., 2005; Ietswaart et al., 2006). Imitation impairments in these patients also dissociate from deficits in the ability to recognize gestures, suggesting that imitation deficits are not due to more general cognitive impairments (Mozaz, 1992). Our work as well as that of others has revealed that patients with apraxia are overly reliant on visual feedback to guide their movements (Howard et al., 2019; Jax et al., 2006; Mutha et al., 2010; Okita et al., 2017). Depriving patients of visual feedback through blindfolding, for example, results in significantly worse imitation performance for apraxic versus non-apraxic individuals (Jax et al., 2006). Blindfolding also reduces the number of error-correction attempts made by individuals with apraxia, as well as the success of those attempts (Howard et al., 2019). Such findings are suggestive of a dependence on a compensatory mechanism in which visual feedback is required to verify the attainment of one’s desired movement goal. This highlights an important puzzle: why can’t patients with apraxia recognize their errors based on proprioceptive feedback alone?

Proprioception has previously been speculated to support the ability to imitate (Meltzoff, 2007; Meltzoff & Moore, 1977; Mitchell, 2002; Rothi et al., 1991). Indeed, it is possible for neurotypical individuals to accurately imitate in the dark or when the movements to be imitated take one’s limbs beyond the field of view (i.e., behind one’s back). One prominent hypothesis suggests that proprioception contributes to imitation through kinesthetic-visual matching, or the ability to translate proprioceptive feedback into an estimate of how the body currently looks (Mitchell, 2002). Note that this hypothesis assumes that the imitation goal exists in a visual format, and that proprioceptive information is transformed into a visual representation as a means of estimating how one’s own body looks in comparison to the visual goal (Fig. 1A). Under this hypothesis, patients with apraxia are poor at imitating when blindfolded because they cannot estimate where their limbs are positioned in visual space (even when using the ipsilesional arm), either due to a primary proprioceptive impairment (but see: Nobusako et al., 2018; Semrau et al., 2013), or due to deficient ability to transform proprioceptive information into a visual format to enable matching with perceived visual information.

**Figure 1:**
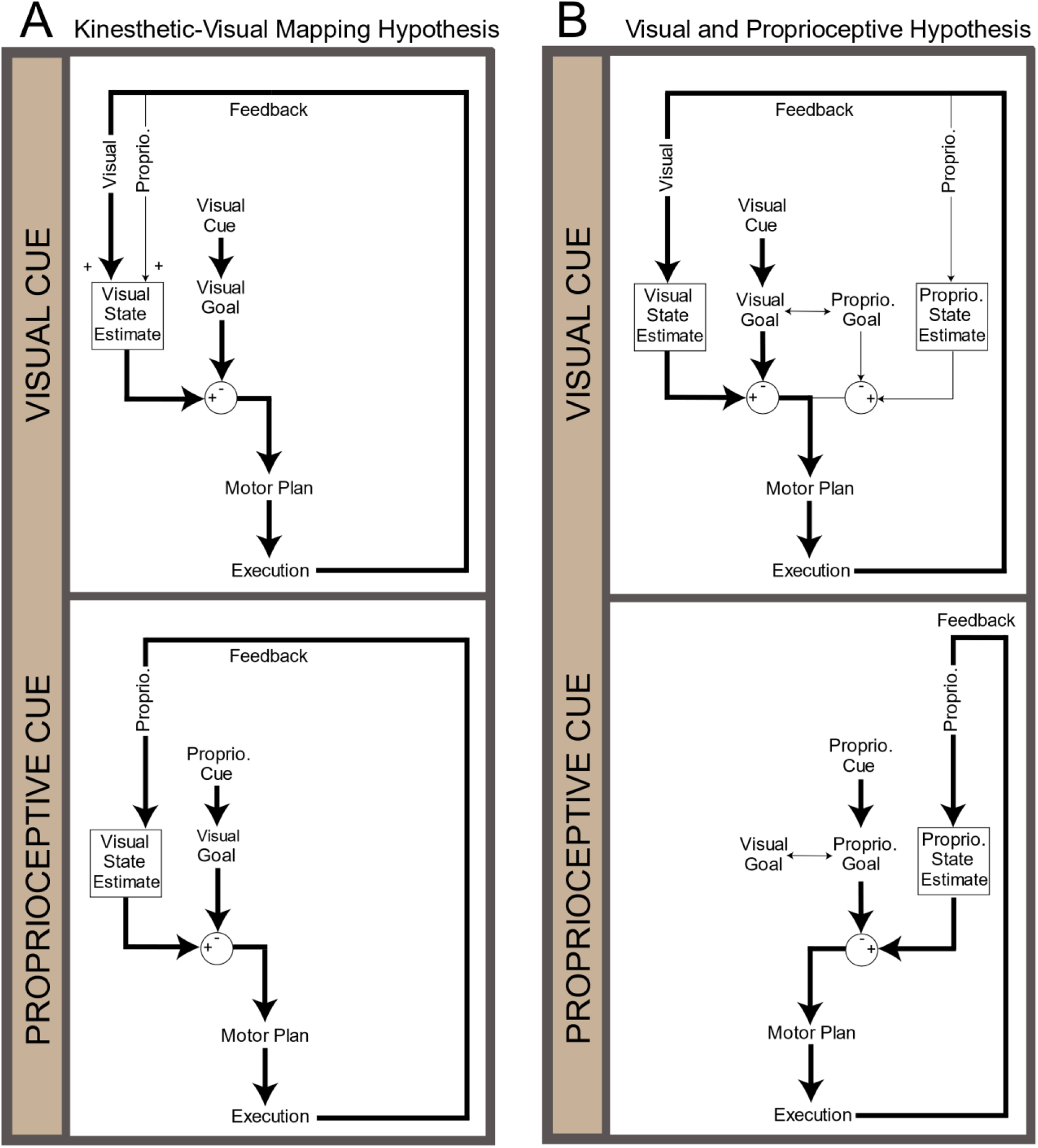
Two alternative hypothesized pathways for imitation. We consider two contrasting hypotheses in the context of our task, where cues are presented purely visually or purely proprioceptively (top vs. bottom). Initial observation of a cue (visual or proprioceptive) leads to the formation of a desired movement goal which is used to plan the movement and supports error corrections. In these diagrams, bold arrows reflect the primary pathway used to guide imitation. A) According to the Kinesthetic-Visual Matching Hypothesis, the observed stimulus (visual or proprioceptive) is used to create a movement goal represented in visual space. During execution, visual and proprioceptive feedback is integrated to form a state estimate of how the body appears visually, which is compared against the visual movement goal to monitor the current movement for errors. B) According to our Visual and Proprioceptive Hypothesis, observation of a movement can produce goals that exist in both visual and proprioceptive forms (i.e., that specify how the action should look and how it should feel), presumably with some interaction between these goal representations. Sensory feedback is primarily compared with its corresponding goal in a modality-specific manner. Thus, in cases where no visual feedback is available, such as is the case in our purely proprioceptive condition (bottom panel, movement is primarily guided according to a goal of how the movement should feel. According to this hypothesis, there is no need to depend upon a representation of how the movement should look.

In contrast, in response to the evidence of the importance of visual feedback in apraxia, we have proposed the Visual and Proprioceptive Hypothesis (Buxbaum & Kalénine, 2014; Howard et al., 2019) where, in the healthy system, movements to be imitated are not only represented visually, but also proprioceptively. That is, we hypothesized that when we imitate, we form proprioception-based movement goals that represent how movements are supposed to feel, which compliment vision-based goals of how movements should look (Fig. 1B). Note that although this hypothesis specifies that visual and proprioceptive goal representations are compared to separate visual and proprioceptive feedback streams, this framing is adopted largely to emphasize the existence and importance of proprioception-based information in the goal representation. Here we remain agnostic as to whether visual and proprioceptive goals are maintained as separate representations or rather integrated into a multimodal representation (and if so, how visual and proprioceptive information is weighted); some evidence suggests that multimodal goal representations are likely (van Beers et al., 2002). Regardless, an emphasis on proprioceptive (in addition to visual) goal representations is consistent with recent work suggesting that we recall proprioceptive memories of movements to recognize and guide future behavior (Kumar et al., 2021; Wong et al., 2012). We further hypothesized that patients with apraxia are impaired at representing proprioceptive goals, leading to the observed overreliance on visual feedback and hence particularly poor performance when imitating while blindfolded. However, we note that this hypothesis may not directly explain all of the patterns of behavior that have been reported in apraxia (e.g., imitating using a mannequin, Goldenberg, 1995).

In this study, we directly tested our hypothesis by asking neurotypical controls and patients with left hemisphere strokes (with a range of apraxia severity) to imitate actions when provided with only proprioceptive information about the desired movement to be copied; individuals were required to reproduce those actions using only proprioceptive feedback with the explicit goal of reproducing how the movements felt. Note that cuing movements proprioceptively and performing them without visual feedback means that participants need not translate proprioceptive feedback into visual estimates of how the limb looks in space in order to imitate. Moreover, evidence suggests that proprioceptively perceived movement goals do not need to be translated into a visual representation: biases when reaching lie in different (sometimes even orthogonal) directions when aiming for visual versus proprioceptive targets (Block & Bastian, 2010; Smeets et al., 2006). Thus, we can be relatively confident that our proprioceptive condition is a fairly pure assay for imitation guided by a representation of how that movement should feel, and is not obligately mediated by an intermediate movement goal representation in a visual format. This proprioceptive cue condition was contrasted against one in which individuals were cued visually by watching a moving cursor (providing no information about desired body configurations) and reproduced the same motion by guiding a cursor with their unseen arms. Performance differences in these two conditions allowed us to assess whether we indeed represent imitation goals proprioceptively. We also examined whether differences in performance could be explained by an individual’s proprioceptive acuity (i.e., a primary proprioceptive deficit). Finally, we examined how performance in these imitation conditions related to two measures often used to assess apraxia severity (Buxbaum et al., 2014b; Kalénine et al., 2010; Tarhan et al., 2015). We hypothesized that relative deficits in the proprioceptive cue condition would be associated with deficient performance on a more conventional test of imitation ability, but not with a test of gesture recognition that probes action semantic knowledge but does not require overt movement planning (van Elk et al., 2014). Together, this work provides insight into how we represent movement goals for imitation and the underlying mechanisms of imitation impairments in patients with apraxia.

## 2 METHODS

### 2.1 Participants

Thirty chronic stroke survivors with a single stroke in the left-hemisphere and no history of other neurological disorders (60.9 ± 10.4 years old; 16 Female) were recruited through the Moss Rehabilitation Research Institute’s Research Registry (Schwartz et al., 2005). All patients were right-handed prior to their stroke. To ensure that patients could understand task instructions, we excluded those that exhibited severely impaired verbal comprehension, classified as a score lower than four out of twenty on the comprehension subtest of the Western Aphasia Battery (Kertesz & Grune & Stratton, 1982). Patients were not specifically selected for presence or severity of apraxia, thus we expected our patient group to exhibit a range of apraxia scores on our laboratory’s apraxia assessments (Buxbaum et al., 2014b; Kalénine et al., 2010; Tarhan et al., 2015). Fourteen age-matched, right-handed neurotypical controls (62.4 ± 7.8 years old; 10 Female) were also recruited through the Research Registry. All participants provided written informed consent prior to study participation in accordance with the guidelines of the Einstein Healthcare Network Institutional Review Board and received compensation for their time. Sample size was chosen to be similar to prior studies examining the kinematics of patients with stroke (e.g., Haaland et al., 1999; Schaefer et al., 2007; Wong et al., 2019). Study procedures and analyses were not pre-registered before they were conducted.

### 2.2 Equipment/Materials

Participants performed all tasks using their left (ipsilesional, less-affected) arm to avoid hemiparesis in the contralesional (right) arm. Note that apraxia, when present, affects both arms. For the Visual/Proprioceptive Imitation Task (“VP Imitation Task”), participants were seated in a robotic exoskeleton (Kinarm Exoskeleton Lab, Kinarm, Kingston, ON, Canada) with their arms supported horizontally just below shoulder height by troughs at the hand, forearm, and upper arm. In cases where impaired arm mobility precluded comfortably resting the paretic (contralesional, right) arm in the troughs, the paretic arm (which did not perform the task) was instead propped on a pillow in the patient’s lap to ensure that the shoulders remained level and centered in the chair throughout the experiment. Participants viewed stimuli presented by an LCD monitor that was reflected in a mirror located directly above their arms, resulting in stimuli that appeared in the plane of the participant’s arms. The center of the workspace was determined according to the position of the index finger when the shoulder angle was 35 degrees from the coronal plane and the elbow angle was 110 degrees. Vision of the arms and torso were occluded by an opaque screen located directly beneath the mirror, as well as a bib extending from the screen, parting just around the participant’s neck, and ending at the chair behind them. This same setup was used to test proprioceptive acuity (see below), with participants reporting their sensed arm displacement using a small 2-axis analog joystick (Adafruit, New York, NY, USA) that was attached to the left-hand trough. All tests using the robotic exoskeleton were programmed in Simulink (The Mathworks, Natick, MA, USA). Finally, a test of visual trajectory-matching ability was performed. Participants were seated at a desk in front of a desktop computer and responded with keyboard presses to stimuli presented on the screen via a program written in C++. All patients also previously completed tests of gesture imitation and recognition (Buxbaum, Kyle, et al., 2005; Tarhan et al., 2015). Code for these tasks can be found on the lab’s Github page (e.g., https://github.com/CML-lab/VisProp_Imitation). Data and analysis code are accessible at https://osf.io/gvpy8/?view_only=24c4dea37dec4c5a858a5e54a3ccb113).

### 2.3 Procedure

#### 2.3.1 VP Imitation Task

In the VP Imitation Task, we examined participants’ abilities to copy meaningless trajectories that were cued either visually by watching a cursor move on the screen, or proprioceptively (without vision) by feeling passive motion of the left arm as guided by the robot (Fig. 2), analogous to previous studies probing the role of visual and proprioceptive feedback (e.g., Adamovich et al., 2001). Here we refer to these conditions as “visual” and “proprioceptive” imitation to emphasize the sensory modality in which the movement to be imitated was cued and for which the dominant source of feedback was likely to be during imitation. Note that testing a truly visual-only imitation task would have required blocking or disrupting proprioceptive feedback of the limb in some way. Here we simply provide cursor-only feedback that includes no information about the desired arm configuration; previous research suggests that neurotypical individuals and (to an even greater degree) patients with left hemisphere stroke rely primarily on visual feedback when it is available (Bagesteiro et al., 2005; Block & Bastian, 2010; Howard et al., 2019; Jax et al., 2006; Mutha et al., 2010; Okita et al., 2017; Over, 1966). Hence our two conditions likely biased individuals to imitate using the sensory modality in which they were cued. Visual and proprioceptive imitation were performed in two separate blocks; block order was counterbalanced across participants within each group (patients, controls).

**Figure 2:**
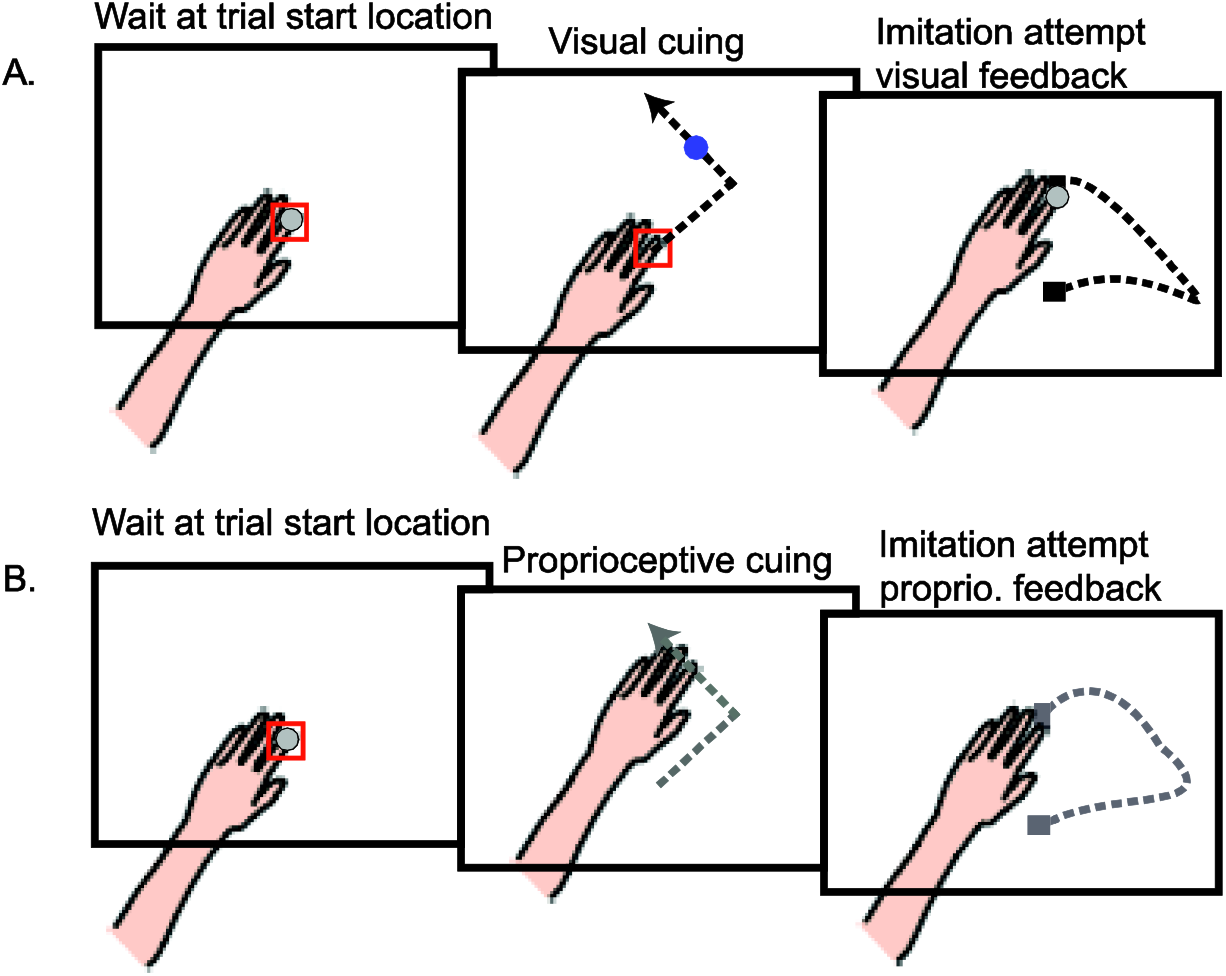
Procedure for different cuing modalities in the VP Imitation Task. Each trial began once the participant reached the start location (red square); in all cases the hand and arm were not visible to the participant. (A) In the visually cued trials, participants held their hand in the start location as they watched a blue dot traverse the desired trajectory. Once the dot disappeared, they imitated the motion by controlling a cursor representing their index finger. (B) In proprioceptively cued trials, the participant’s left hand was moved by the robot along the desired trajectory. Upon completion of the trajectory they were moved back to the start location, whereupon participants imitated the cued trajectory. No visual cursor feedback was given during cuing or execution in these proprioceptively cued trials.

Movement trajectories were composed of one to three straight or semicircular segments that were specified as Bezier curves with bell-shaped velocity profiles. Trajectories were subjectively spatially balanced across the workspace; together, all movements covered as much of the workspace as possible. Each segment was 2 seconds in duration and traversed approximately 9 cm. In each block of 48 trials, participants first completed four repetitions of four single-segment “easy” movements, pseudorandomized such that participants saw each of the four easy trajectories in a random order before any trajectory was repeated. Participants then completed four repetitions of eight “hard” two- or three-segment movements, which were also pseudorandomized such that all eight trajectories were presented before any were repeated. Because patients with apraxia are relatively unimpaired when they can specify movements as point-to-point reaches to a target goal (Buxbaum et al., 2005; Ietswaart et al., 2006), and because patients with apraxia are known to be most impaired when asked to perform heterogeneous movement sequences (Harrington & Haaland, 1992), we expected patients to be relatively more impaired on hard versus easy trials compared to controls (i.e., a group by difficulty interaction). This prediction is analogous to what we observed previously (Wong et al., 2019). Because of this potential alternative strategy for performing the easy movements, however, we opted to focus on analyzing the hard trials in the majority of our analyses. To equate trajectory difficulty across the visual and proprioceptive imitation blocks while preventing participants from performing the same set of movements in the two blocks, movements were mirrored about the midline in the two cuing conditions, with the specific subset of mirrored movements assigned to the visual and proprioceptive condition counterbalanced across participants. In all cases, participants received no explicit feedback about their imitation accuracy aside from what they could observe of their own behavior.

In the visual imitation block, the participant was given feedback of their hand location throughout the block in the form of a cursor (1.5 cm diameter) projected on the tip of the index finger. The participant was instructed to position the cursor inside a small red box at the start position and keep it there while watching a blue dot (1.5 cm diameter) traverse the screen in the desired trajectory. The dot left no trace of its passage as it moved. Participants were instructed to pay close attention to the dot and remember the trajectory along which it moved. At the end of the trajectory, the blue dot disappeared. After 500 ms, an auditory tone prompted the participant to begin imitating the trajectory. Once the experimenter confirmed that the participant had completed their movement, the experiment was manually advanced to the next trial. In the proprioceptively cued block, the robot initiated each trial by guiding the participant’s arm to the start box. The participant was instructed to keep their arm relaxed throughout. After 800 ms, a tone sounded, and the robot guided the arm along the desired trajectory. Participants were instructed to pay close attention and remember how this movement felt so that they could imitate it after the movement was over. At the conclusion of the movement, a second tone sounded to indicate that the movement was complete, and following a brief pause the robot returned the hand to the start position and released it, requiring the participant to maintain their unseen hand within the start box. After 500 ms, a start tone indicated that the participant should begin imitating the movement trajectory. While moving, the participant received no visual feedback of their hand position, encouraging them to reproduce how the movement felt based on proprioceptive feedback of the limb. Once the participant stopped moving, the experimenter manually progressed to the next trial.

In designing this task, a question arose as to whether previous experience in seeing the same movements cued visually could improve proprioceptively-cued imitation, in the same way that action observation modifies future movement planning and behavior (Gonzalez-Rosa et al., 2015; Stefan et al., 2005). As the VP Imitation Task was designed to specifically avoid potential repetition effects, to test for a cuing-order effect a subset of participants (N = 22 patients, 60.2 ± 11.0 years old, 10 female; N = 10 controls, 61.3 ± 8.5 years old, 6 female) also completed two additional blocks. In these blocks, participants imitated four additional two- and three-segment trajectories that were not mirror-inverted between the two cuing conditions. Each trajectory was performed twice in randomized order within each cuing condition, and participants always experienced the visual cuing condition first, followed by the proprioceptive cuing condition. Note that this manipulation also gives us insight into whether movements in the proprioceptively cued condition must be transformed into a vision-based goal representation; if so, we would expect that priming of such a representation (i.e., by previously observing the movement visually) would improve performance in the proprioceptive imitation condition.

#### 2.3.2 Control task: Proprioceptive Acuity

Immediately following the VP Imitation Task, participants completed a test of proprioceptive acuity (modeled after Wilson, Gribble, Wong 2010). For this test, the robot first positioned the participant’s left arm at a reference position, which remained fixed throughout the entire experiment. After a tone and a 1750 ms delay, the robot then moved the arm along a distracting, curved trajectory (created from a set of 2-3 Bezier curves) in the horizontal plane before stopping directly to the left, to the right, nearer, or farther from the start position. Participants were informed that the endpoint would always be in one of five locations: one of the four displaced positions noted above or at the same point as the start position. When the arm stopped moving, participants were asked to indicate whether the start and end arm positions were the same. Although participants were not aware of it, the choice of “same” was never correct; the closest the hand was ever placed relative to the reference position was 0.5 cm. Participants responded by verbally indicating whether the final position was different from the reference position or not (“yes” or “no”). On each trial, the distance from the reference position was determined using a separate staircase for each of the four displaced directions. Two staircases were initialized at a hand displacement of 0.5 cm from the start position, and the remaining two were initialized at 10 cm. If participants correctly detected a displacement, the displacement decreased by 0.5 cm for the next trial; if they responded incorrectly the displacement increased by 0.5 cm. Once a given staircase switched from decreasing to increasing 3 times within 8 trials, we considered that staircase to have converged to a detection threshold and it was not tested further. The task was finished when participants reached a threshold or completed 25 trials in all four directions, whichever came first.

#### 2.3.3 Control task: Visual Recognition

Finally, to test whether any deficits in the VP Imitation Task were attributable to impairments in perceiving and remembering trajectories, participants completed a two-alternative forced choice test of visual trajectory recognition and working memory. This task was analogous to the visual block in the VP Imitation Task in terms of the cursor movements presented and the stimulus timing. With participants seated in a chair in front of a computer, they saw a black 1 cm dot move in one of the same trajectories as encountered during the VP Imitation Task; as before, the dot left no trace of its passage, so participants had to remember its motion path. After the movement was completed, there was a 500 ms delay and then two static images of trajectories appeared vertically along the screen’s midline, one colored blue and one colored red. Participants had to report which of the two trajectories matched the motion they had just watched by pressing a corresponding red (x key) or blue (z key) button on the keyboard.

#### 2.3.4 Apraxia Battery

Prior to participation in this study, all patients previously completed our laboratory’s tests of the imitation of meaningless gestures and the ability to recognize gestures. Because it lacks overt motor planning requirements and instead probes knowledge of action semantics (i.e., knowledge of the actions associated with objects; van Elk et al., 2014), the latter test served as a negative control as it was not expected to relate to performance on the main experimental task.

In the test of meaningless gesture imitation (Buxbaum et al., 2014), participants watched 14 videos of novel and semantically meaningless gestures. On each trial, participants were shown the gestures twice in succession and were asked to imitate the second occurrence of each gesture as accurately as possible using the left (ipsilesional) hand. Imitation attempts were video recorded, and trained and statistically reliable coders scored imitation attempts as correct or incorrect on four dimensions (hand posture, arm posture, amplitude, and timing) (Buxbaum et al., 2000). An average of these four scores was used as a measure of performance accuracy for each gesture; trial scores were averaged together to yield a measure of meaningless gesture imitation ability for each participant.

In the test of gesture recognition (Kalénine et al., 2010; Tarhan et al., 2015), participants performed a two-alternative forced-choice task. On each of 24 trials, an action name (e.g., “combing hair”) was presented, followed by two videos of actions being performed. One video showed the correct action, and the other showed a semantically-related action (e.g., brushing teeth). Participants were asked to choose which of the two videos corresponded with the cued action phrase. A verb-picture matching pre-test (match a picture of a comb to the word “combing”) was also completed to ensure that patients comprehended the verbs that were used in the main task. If patients failed to match a verb to the correct tool in the pre-test, the corresponding action phrase was excluded from the main task. Gesture recognition performance was measured as the adjusted percent of correct trials. This test of gesture recognition relates to another common deficit observed in apraxia, impairment of tool-use abilities; hence this test allowed us to examine whether our findings reflect apraxia severity more broadly.

### 2.4 Statistical Analysis

#### 2.4.1 VP Imitation Task

For the VP Imitation Task, motion of the left fingertip was recorded at 1000 Hz and analyzed in MATLAB using custom scripts. Recorded data (position, velocity) were smoothed with a second-order Savitzky-Golay filter with a frame size of 19 samples. Movement start and end were identified as the times when hand velocity exceeded or fell below 0.04 m/s respectively and were verified by visual inspection. Movements were time-normalized and imitation accuracy was quantified using a Procrustes Distance (PD) metric (Goodall, 1991). This measure quantifies the remaining dissimilarity between two trajectories following a series of affine transformations (translation, rotation, and scaling) that aligns the trajectories to the best possible extent (Fig. 3). Using this algorithm, we described the difference between the participant’s movements and the desired trajectory (i.e., the motion of the cursor/robot; larger PD values indicate greater dissimilarity or worse performance accuracy). While PD is typically bounded between 0 and 1 (Gower, 1975), the upper bound is relaxed when the trajectory is allowed to be scaled differently along each dimension of the data (Rohlf & Slice, 1990). We used this modified PD metric in our analyses since allowing the trajectory to scale differently along the X and Y dimensions greatly improved the quality of the algorithm’s fits as noted by careful visual inspection of a subset of our dataset.

**Figure 3:**
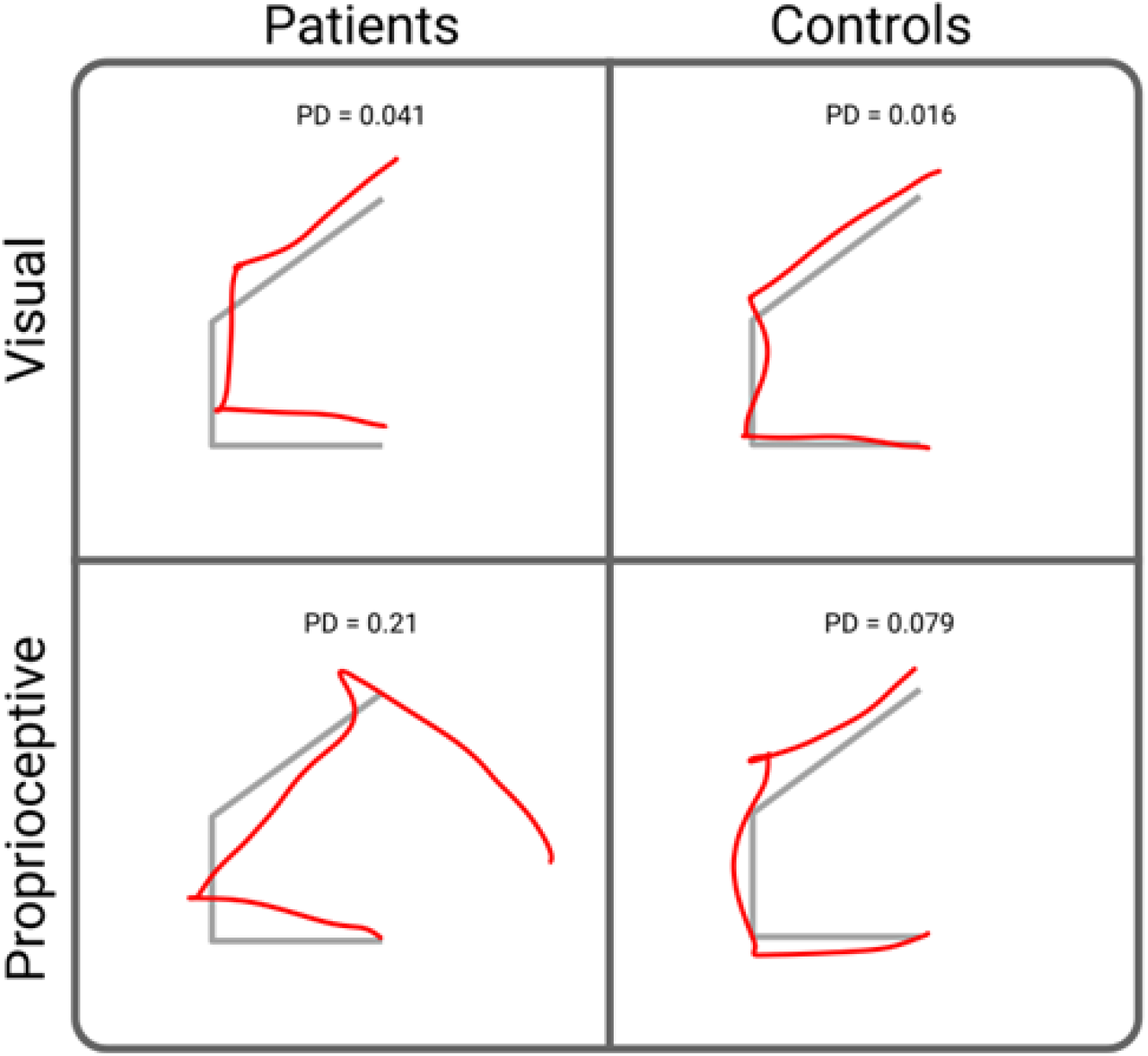
Imitation performance as quantified by the Procrustes Distance (PD) algorithm. Example of raw trajectories produced by a representative patient and control participant in the visual-imitation and proprioceptive-imitation conditions are shown in red, with the desired trajectory in gray beneath it. Scores for each example trajectory are given at the top of each panel. Raw trajectories are translated, scaled, and rotated until they align with the desired trajectory as best as possible; dissimilarity is then quantified as the PD score, or the remaining sum of squared differences of the coordinates between each shape. Higher PD scores reflect less accurate imitation.

Even if individuals imitated with low accuracy, this does not indicate whether the errors were produced with any degree of consistency. If individuals were consistent in their behavior, we would expect to see systematic errors on each repetition of the same movement; hence, cross-trial variance should remain roughly similar from the beginning to the end of a given movement path. In contrast, if individuals make inconsistent errors each time they attempt to reproduce a given movement path, their movements should diverge over time and we would expect to see cross-trial variance dramatically increase as the movement progresses. Thus, to quantify consistency, we examined the change in variance across the movement on a per-trajectory, per-cue basis for each participant. We used the change in variance from the start to the end of the movement as a measure of inter-trial consistency. To quantify change in variance, we first aligned the traces for all four repetitions of a given trajectory to movement start and found the position of the hand in Cartesian coordinates at the time of the first velocity peak (i.e., at the first local maximum in the magnitude of the velocity trace). Total Variation early in the movement was computed as the trace of the covariance matrix of hand position at the first velocity peak. Analogously, we computed Total Variation late in the movement by aligning the same four trajectories to the end of the movement and computed the trace of the covariance matrix of hand position at the last velocity peak before movement offset. We realigned the data at movement end to reduce the effect of noise accumulation throughout the movement as well as to reduce the interdependence of variance estimated at the two time points (yielding a more conservative estimate of the change in variance across the movement). Finally, we computed the change in variance throughout the movement, ΔTV, as the Total Variation late in the movement minus Total Variation early in the movement.

We also examined temporal measures during the VP Imitation Task. Reaction time was calculated as the time from the go cue until participants initiated their movement. Movement time was calculated as the time from movement start to movement end.

Finally, we assessed whether prior experience at imitating a trajectory visually provides a benefit when imitating the same trajectory proprioceptively. To examine this, we computed the difference in average PD over each trajectory when imitating a trajectory proprioceptively versus visually. Only “hard” two- and three-segment movements were assessed. We examined the difference in performance when individuals experienced the exact same trajectory in both conditions (repeated trials) in close succession, compared to the difference in the standard VP Imitation Task when individuals experienced the mirror-image version in the two cuing conditions (non-repeated trials). The average latency between presentation of the same trajectory in the visual and proprioceptive conditions was 74 seconds (range 48-100 secs).

For all metrics above, statistical analyses were performed in R (R Core Team, 2020). using the *lme4* package (Bates et al., 2014) to fit mixed effect models to our behavioral data. Significant effects were identified using likelihood ratio tests comparing models with and without the effect of interest. Specifically, based on our *a priori* hypotheses we examined fixed effects of group, difficulty, and cue type, along with two-way interactions between group (patients, controls) and difficulty and between group and cue type. When appropriate, we included random effects of participant and trajectory. In all models, we also checked for effects of order and performance on our two control tasks. Our *a priori* hypothesis was that neither order nor the control tasks would significantly affect our data; thus, we only retained these terms in our models when they were found to be significant. Since PD is a bounded metric and RT is known to not be normally distributed, we applied a logarithmic link function when analyzing models involving these dependent variables. When an interaction was found to be significant, we performed post-hoc analyses to examine the interaction by dividing the data along one factor and examining the effect of the second factor separately in each subset of the data. Holm-Bonferroni corrections were applied to correct for multiple comparisons.

#### 2.4.2 Control Tasks

For each participant, proprioceptive acuity in the control task was calculated by fitting a mixture model to response data (correct detection of arm displacement on each trial as a binary outcome measure, versus the displacement size on that trial in cm) using Expectation-Maximization. This mixture model consisted of a logistic sigmoid function describing arm displacement detection plus a uniform distribution that was used to identify outliers. We took the inflection point of the sigmoid, representing the distance at which the participant correctly detected a displacement on 50% of trials, as an estimate of proprioceptive acuity. Proprioceptive acuity was examined as a potential fixed effect in all mixed effect models above. We also directly correlated proprioceptive acuity with imitation accuracy from the main task.

In the visual trajectory-matching control task, we calculated the proportion of correct trials as the sum of correct trials divided by the total number of trials in the task. This proportion was examined as a potential fixed effect in the mixed effects models above, as well as directly correlated with imitation accuracy from the main task.

#### 2.4.3 Apraxia Scores

To test the prediction that disproportionate impairments in proprioceptive imitation would be related to limb apraxia on a task requiring movement production, we computed the residuals of the relationship between proprioceptive and visual imitation accuracy on the experimental tasks according to our PD metric above. We correlated these residuals with performance on our laboratory’s well-studied measure of imitation of meaningless movements (Buxbaum et al., 2014). As deficits in proprioceptive imitation were hypothesized to reflect specific deficits in proprioceptively-related movement planning rather than other aspects of the apraxia syndrome (e.g., deficient gesture knowledge), we also tested the prediction that the correlation between these residuals and a test of gesture recognition should be relatively weak (Kalénine et al., 2010).

## 3 RESULTS

### 3.1 Stroke Patients Were Disproportionally Inaccurate in Proprioceptive Imitation

Our primary hypotheses were that imitation ability would differ when movements were cued visually or proprioceptively, and that this difference would be exaggerated in patients with left-hemisphere stroke compared to neurotypical controls. We used a Procrustes distance (PD) metric to assess imitation accuracy (Fig. 4). Performance in the visual and proprioceptive imitation conditions was correlated across all participants (*r²*(42) = 0.35, *p*<.001), driven largely by the patient group (patients: *r²*(28) = 0.33, *p*<0.001; controls: *r²*(12) = −0.08, *p* = 0.97), likely because controls were more uniformly accurate. We next conducted a mixed-effects model to examine imitation ability across groups and conditions. We first tested the effects of cuing-condition order and the control tasks on imitation accuracy. Neither Order (χ^2^(1) = 0.35, *p* = 0.55) nor proprioceptive acuity as measured in our control task (χ^2^(1) = 0.21, *p* = 0.65) significantly affected imitation accuracy, so these terms were removed from the model. Visual trajectory recognition had a significant effect on imitation accuracy (χ^2^(1) = 4.17, *p* = 0.041), so this term was retained in the model as a nuisance parameter. Using this model, we observed a significant main effect of group (χ^2^(3) = 24, *p*<0.001)^1^. We also observed a significant main effect of cue (χ^2^(2) = 181.10, *p*<0.001), indicating that people were less accurate at imitating proprioceptively-cued movements compared to visually cued movements (Fig. 4A). Importantly, there was a significant interaction between group and cue (χ^2^(1) = 9.26, *p*<0.01), supporting our hypothesis that compared to controls, patients were disproportionately worse in the proprioceptive versus visual cue condition (points falling farther above the unity line in Fig. 4B; difference between imitating visually and proprioceptively cued movements: patients, 0.14 ± 0.11; controls, 0.028 ± 0.087). Not surprisingly, we additionally observed a significant effect of difficulty, indicating that hard trajectories (i.e., trajectories with more segments) were indeed more challenging to imitate for both groups (χ^2^(2) = 8.43, *p* = 0.030). However, counter to our hypothesis that patients would be particularly impaired at hard trajectories because they could adopt an endpoint-aiming strategy in easy trials rather than imitating the movement¹ there was no significant interaction between group and difficulty (χ^2^(1) = 0.025, *p* = 0.87). This suggests that both patients and controls responded similarly to the difficulty manipulation. One possible reason for this finding could be that both groups opted for an aiming strategy on easy trials, and thus the two difficulty levels reflect a change in movement strategy rather than a manipulation of imitation difficulty. To avoid any such potential confounds, for the remainder of the analyses we focused on performance in the hard condition. Nevertheless, overall our findings suggested that patients were often disproportionately worse at imitating proprioceptive cues than visual cues compared to neurotypical controls.

**Figure 4:**
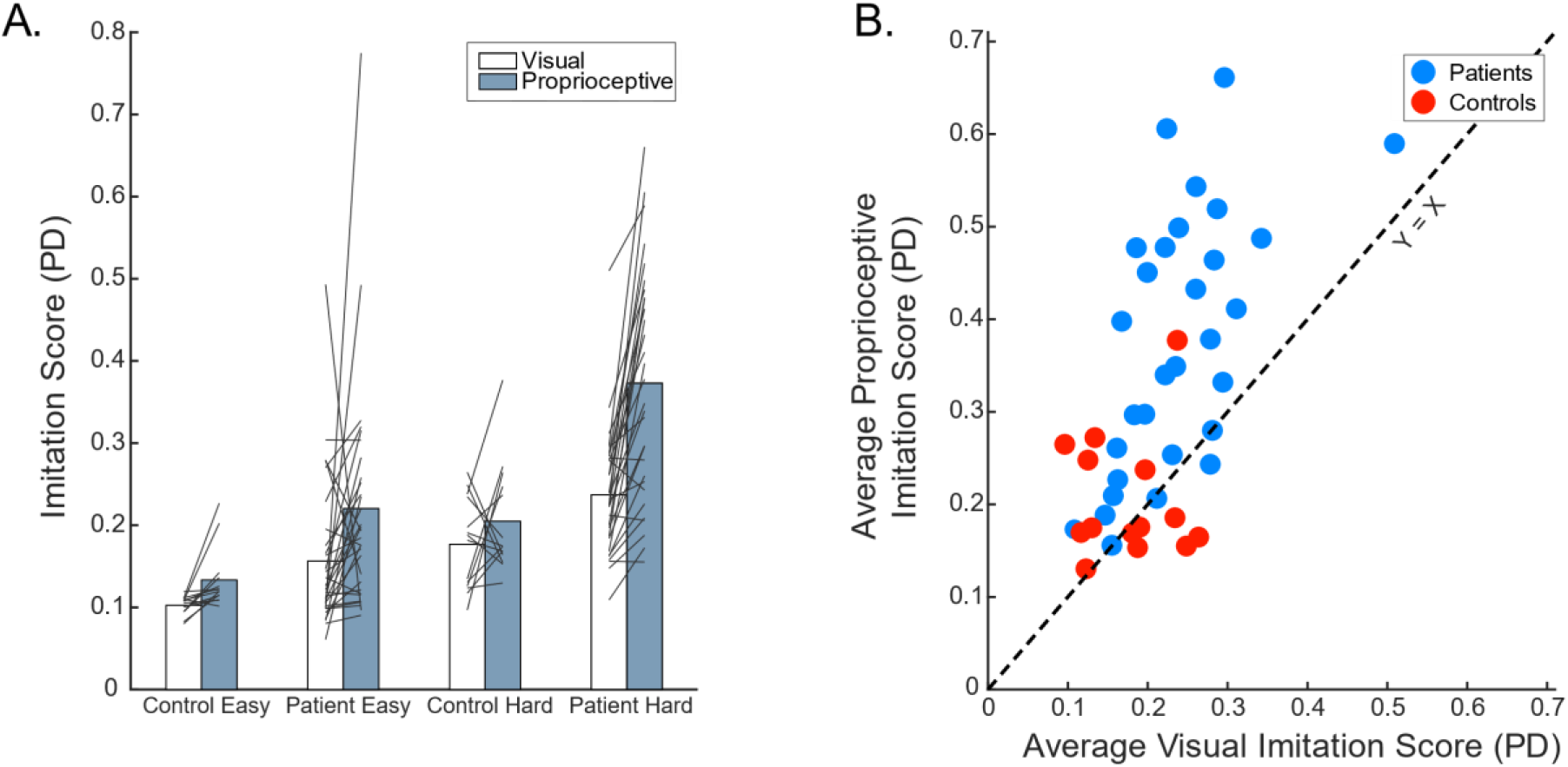
Accuracy in the VP Imitation Task. (A) Imitation performance (Procrustes dissimilarity, PD) is plotted for all individuals across all task conditions (larger scores are worse). In general, patient performance tends to be worse than controls, particularly when cued proprioceptively (grey versus white bars) or when the movements are more difficult (i.e., “hard” multi-segmented versus “easy” single-segmented trajectories). Grey lines reflect individual subjects’ scores in the visual and proprioceptive conditions. (B) Relationship between visual and proprioceptive imitation in the hard condition, compared to the equality line (y = x). While most participants exhibit slightly worse imitation performance when cued proprioceptively compared to visually, patients tend to lie farther above the equality line, suggesting a disproportionate impairment in proprioceptive imitation.

One *a priori* question we had was whether imitation performance in the visual or proprioceptive cuing conditions could be attributable to underlying perceptual or memory deficits in a modality-specific manner. In line with our findings above, we noted a significant relationship between visual imitation accuracy and visual recognition score from our visual control task across all participants (Fig. 5A; *r²*(39) = 0.52, *p* < .001), and for patients alone (*r²*(29) = 0.63, *p* < 0.001), but not for controls (*r²*(13) = 0.004, *p* = 0.82). We did not see a significant relationship between overall proprioceptive imitation accuracy and proprioceptive acuity as measured by our proprioceptive control task (Fig. 5B; *r²*(37) = 0.012, *p* = 0.51). Because our findings revealed that patients exhibit *disproportionately* poor performance in the proprioceptive imitation condition compared to the visual condition, we also examined as a *post hoc* test if this disproportionate deficit (i.e., the residuals of the regression between proprioceptive and visual imitation accuracy) could be explained by an underlying proprioceptive deficit. Interestingly, we did not observe a significant correlation between disproportionately poor proprioceptive imitation performance and proprioceptive acuity from our control task (*r²*(37) = −0.013, *p* = 0.48; Fig. 5C), again suggesting that our findings were unlikely to be explained by underlying low-level proprioceptive impairments.

**Figure 5:**
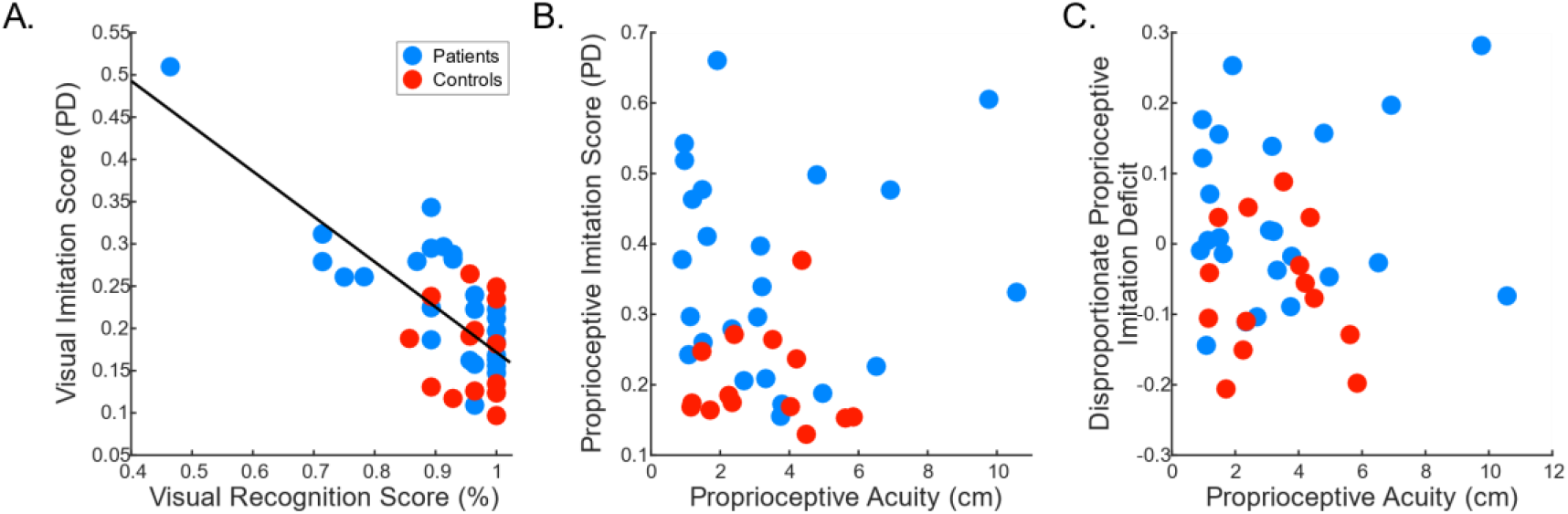
Imitation performance compared to control tasks. (A) Visual imitation ability (average performance in the hard condition) was correlated to the visual recognition control task (*r²*(39) = 0.52, *p* < .001). Note, this correlation persisted even when the individual with a particularly low visual recognition score was removed. This suggests that performance in the visual imitation condition may partially be attributed to the ability to visualize the desired movement shape or hold that shape in memory before responding. (B) In contrast, while 4 patients were outside of the control range in proprioceptive acuity of the ipsilesional arm, no relationship was observed between average proprioceptive imitation scores in the hard condition and proprioceptive acuity. (C) Additionally, no significant relationship was observed between proprioceptive acuity and disproportionate proprioceptive imitation deficits in our experiment, indicating that proprioceptive acuity is unlikely to explain why patients are particularly poor at imitating proprioceptively cued movements.

### 3.2 Repeating Trajectories does not Change Performance

An underlying hypothesis is that if movements are cued proprioceptively and only proprioceptive feedback is available, movement goals can be represented proprioceptively and do not require mediation through a visual representation of what the movement should look like. If instead individuals always rely on a visual goal representation, it stands to reason that recent exposure to the same goal representation evoked visually might confer a performance benefit when imitating proprioceptively (e.g., by helping individuals better visualize how their arm is supposed to move). That is, rather than having to rely on a noisy proprioceptive signal to create a new movement goal representation, if the goals in both imitation conditions are represented visually, individuals should instead be able to recall and reuse a recently formed visually-cued goal representation to support their performance in the proprioceptive imitation condition. To test this hypothesis, we asked whether imitating a trajectory that was cued proprioceptively could be improved if the same movement was previously observed visually. Although we did not detect an effect of order in our analyses, we wanted to test this question directly. Thus, in a subset of participants, we compared performance on the VP Imitation Task against a second pair of blocks in which the trajectories were not mirror-inverted between the two cuing conditions and were always presented first visually then proprioceptively. We observed no significant difference in imitation accuracy as a function of whether the same trajectories had been previously observed visually or not (χ^2^(2) = 1.15, *p* = 0.56); we also did not see a significant difference between groups (although patients tended to perform a little worse than controls in general; χ^2^(2) = 5.37, *p* = 0.067) or a significant interaction (χ^2^(1) = 0.70, *p* = 0.40). Thus, prior experience with having seen a movement was unlikely to provide a benefit when imitating the same movement proprioceptively. This finding, while not providing definitive evidence in support of the Visual and Proprioceptive Hypothesis, is nevertheless suggestive of the notion that the movement goal representation in the proprioceptive imitation condition may not be the same as that evoked in the visual imitation condition.

### 3.3 Proprioceptive Imitation Impairments are Related to a Measure of Praxis Imitation

Our second main hypothesis was that disproportionate deficits in proprioceptively-cued imitation would be related to apraxia severity as assessed by a more conventional imitation task (the imitation of meaningless gestures following visual observation of an actor; (Buxbaum et al., 2014), which provides no proprioceptive information about the desired movement goal. Interestingly, we observed a significant correlation between meaningless gesture imitation and disproportionate impairments in proprioceptive imitation in our VP Imitation Task (i.e., residuals of the proprioceptively cued versus visually cued imitation, Fig. 6A; *r²*(28) = −0.42, *p* = 0.028). Neither visual nor proprioceptive imitation ability alone as quantified in our primary task were found to correlate with performance in the meaningless imitation task (proprioceptive imitation scores, *r²*(28) = −0.30, *p* = 0.11; visual imitation scores, *r²*(28) = 0.057, *p* = 0.76). Meaningless imitation task scores were also unrelated to our control task measuring proprioceptive acuity (*r²*(25) = 0.019, *p* = 0.51). To see if the impairments observed in our task related to other commonly observed apraxic deficits, we also examined whether there was a relationship of the same residuals to a second frequently used apraxia measure, the recognition of meaningful tool-use gestures (known to dissociate from meaningless gesture imitation). As predicted, we observed a non-significant correlation (*r²*(27) = 0.033, *p* = 0.87), suggesting that the observed deficits are not merely a function of overall apraxia syndrome severity but are instead specific to imitation (Fig. 6B). Relatedly, we asked if observed impairments could reflect stroke severity more broadly by testing for a relationship between our findings and stroke lesion volume; again, we observed no such correlation (*r²*(25) = 0.00018, *p* = 0.95). Together, these data suggest that general imitation deficits in apraxia (but not impairments in gesture knowledge) are in part related to the inability to represent proprioceptive movement goals, but are unrelated to primary proprioceptive impairments in estimating limb position.

**Figure 6:**
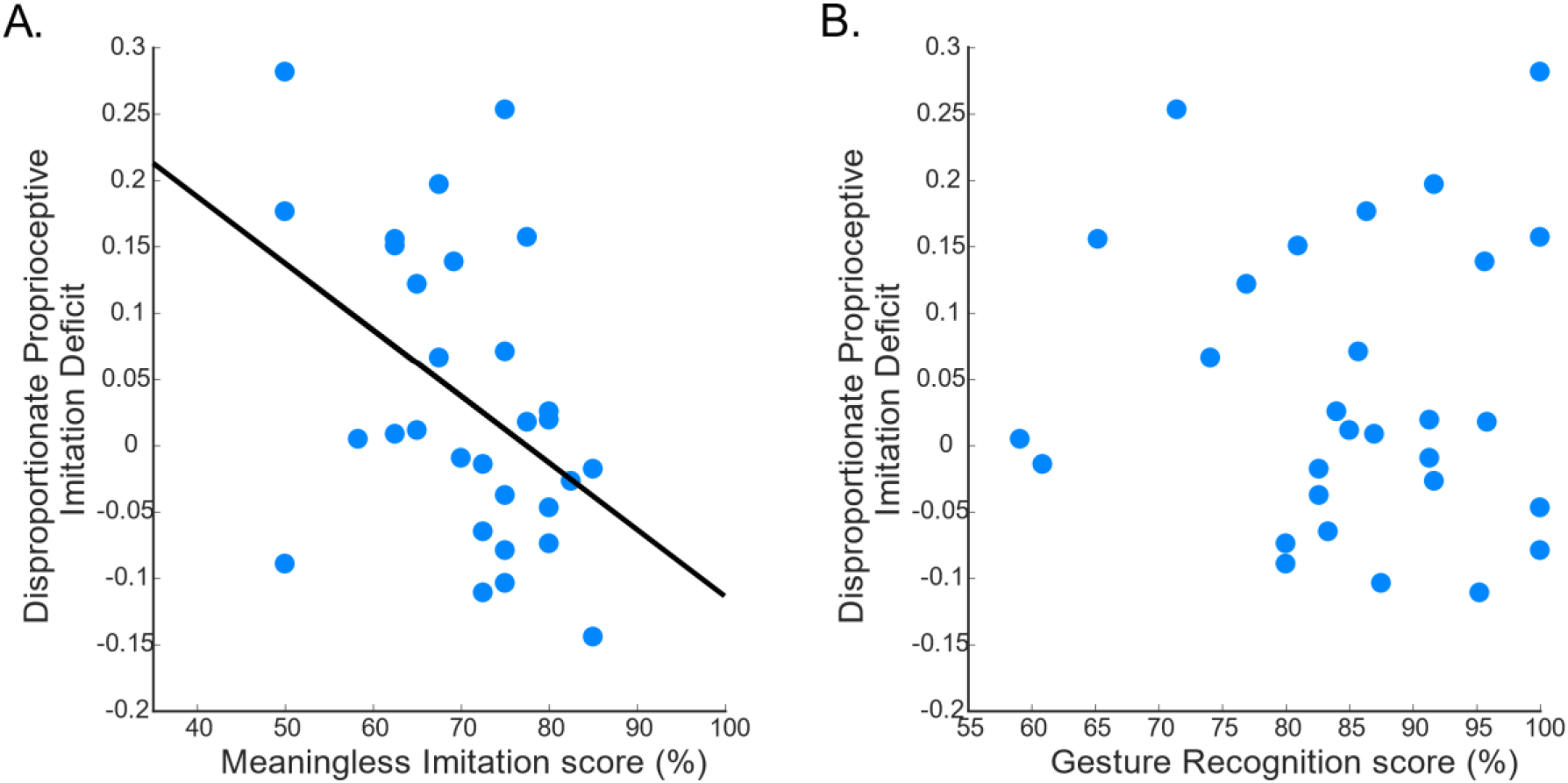
Relationship between our laboratory’s well-studied measures of apraxia and disproportionate proprioceptive-imitation deficits. Disproportionate proprioceptive-imitation deficits were measured as the residuals of the regression between proprioceptive and visual imitation in the hard condition. (A) Disproportionate proprioceptive imitation deficits correlated with imitation of meaningless gestures cued by visual observation of an actor. (B) In contrast, disproportionate proprioceptive imitation deficits were not correlated with a test of recognition of meaningful tool-use gestures.

### 3.4 Patients Imitate Less Consistently

Up to this point, it is unclear whether the imitation errors produced by patients were consistent, as might be expected if, for example, patients had systematic biases in their proprioceptive position sense across the workspace (Fuentes & Bastian, 2010). Thus, as a *post hoc* analysis, we examined how consistent individuals were at imitating movements when cued visually versus proprioceptively by measuring the change in variance from the start to the end of a movement. That is, if individuals made consistent errors from one trial to the next when performing a given movement path, we would expect to see little increase in variance from the start to the end of the movement path; in contrast, if errors were inconsistent, variance should increase considerably from the start to the end of the movement path.

We found that variance growth throughout the movement (as measured by the change in Total Variation, see Methods) increased more during the imitation of proprioceptively-cued movements compared to visually-cued movements (χ^2^(2) = 58.17, *p*<.001). We did not observe a significant effect of group (χ^2^(2) = 0.12, *p* = 0.37), nor an interaction between group and cuing type (χ^2^(1) = 0.26, *p* = 0.51), suggesting that on average, patients and controls exhibited similar increases in variance within a given movement path.

When more closely examining the difference in variance growth for movements cued proprioceptively versus visually (Fig. 7A), we again noted some individuals, particularly patients, who exhibited disproportionately less consistent behavior when movements were cued proprioceptively. We therefore assessed whether the individuals who exhibited disproportionately worse imitation accuracy when cued proprioceptively were also the same individuals who were much more variable when cued proprioceptively. We observed that the relationship between disproportionate proprioceptive imitation accuracy (the residuals of the regression between proprioceptive imitation accuracy and visual imitation accuracy) was significantly correlated with the difference in variance growth between proprioceptive and visual imitation in patients (Fig. 7B; *r²*(29) = 0.17, *p* = 0.024), but not in controls (*r²*(13) = 0.014, *p* = 0.20). This correlation suggests that individuals who are particularly impaired at accurately imitating proprioceptively cued movements are also less consistent in their behavior from trial to trial. That is, individuals are not producing systematic imitation errors; imitation impairments affect both bias and variance.

**Figure 7:**
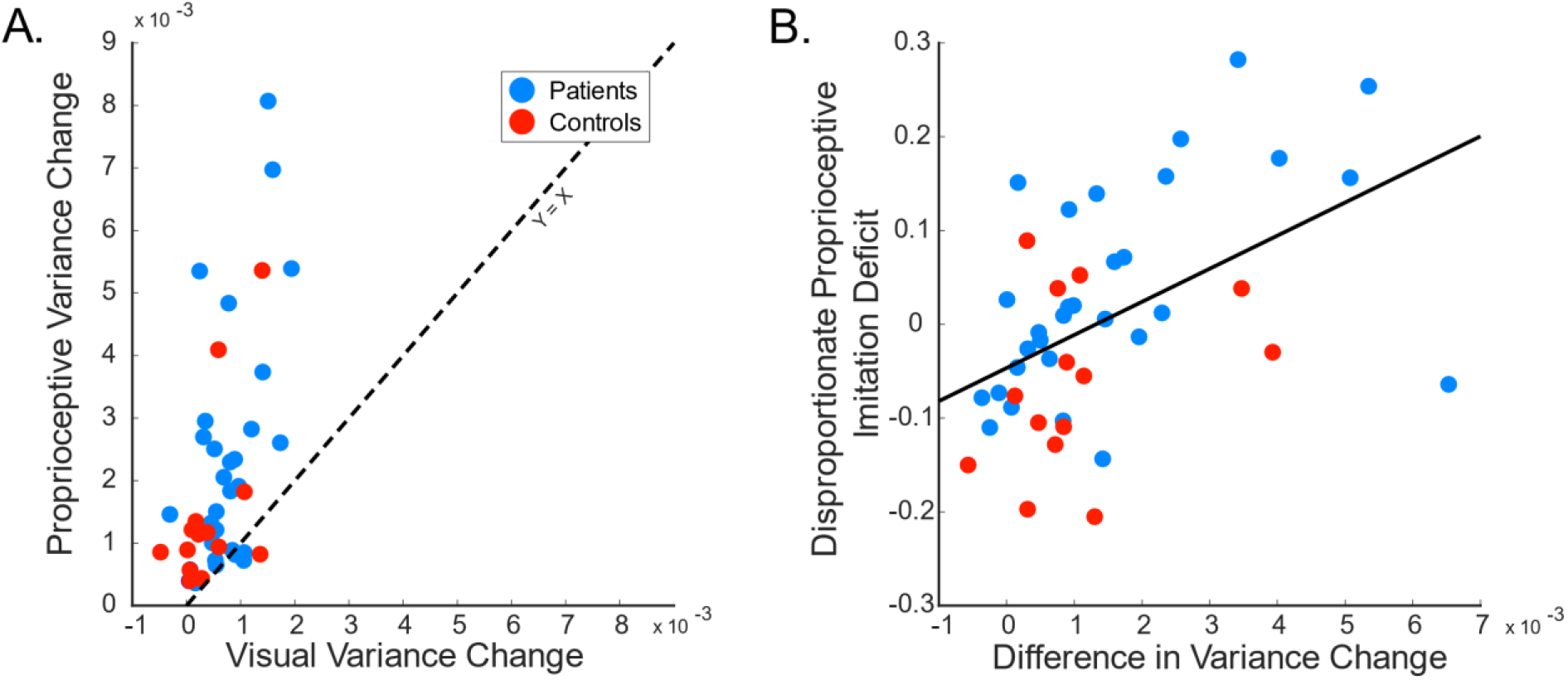
Imitation consistency relates to imitation accuracy. (A) Variance growth across a movement increased much more for proprioceptively cued movements compared to visually cued movements in the hard condition. (B) A significant correlation was observed between disproportionate proprioceptive-imitation accuracy and the difference in variance growth between proprioceptive and visual trials. This suggests that the same individuals are likely to be both disproportionately less accurate and more variable in the proprioceptive imitation condition.

### 3.5 Imitation Impairments Do Not Reflect a Speed-Accuracy Trade-off

As participants were found to be both less accurate and more variable when imitating proprioceptively-cued actions, one concern was that perhaps individuals were exhibiting a speed-accuracy trade-off. Thus, we examined the time it took participants to complete their movements. We observed that movements were actually slower when imitating proprioceptively cued actions compared to visually cued actions (χ^2^(2) = 51.00, *p*<.001). Indeed, if anything we observed a slightly positive relationship between disproportionate proprioceptive imitation accuracy and the difference between proprioceptive and visual movement times (χ^2^(2) = 7.96, *p* = 0.076): individuals who were less accurate at imitating proprioceptively cued actions also tended to move more slowly in the proprioceptive imitation condition (Fig. 8A). We also failed to observe a significant relationship between the difference in movement consistency (i.e., the difference between the change in Total Variation in proprioceptively-cued and visually-cued imitation) and the difference in movement time (χ^2^(2) = 1.07, *p* = 0.59; Fig. 8B). Together, these results suggest that our findings cannot be explained by a speed-accuracy trade-off.

**Figure 8:**
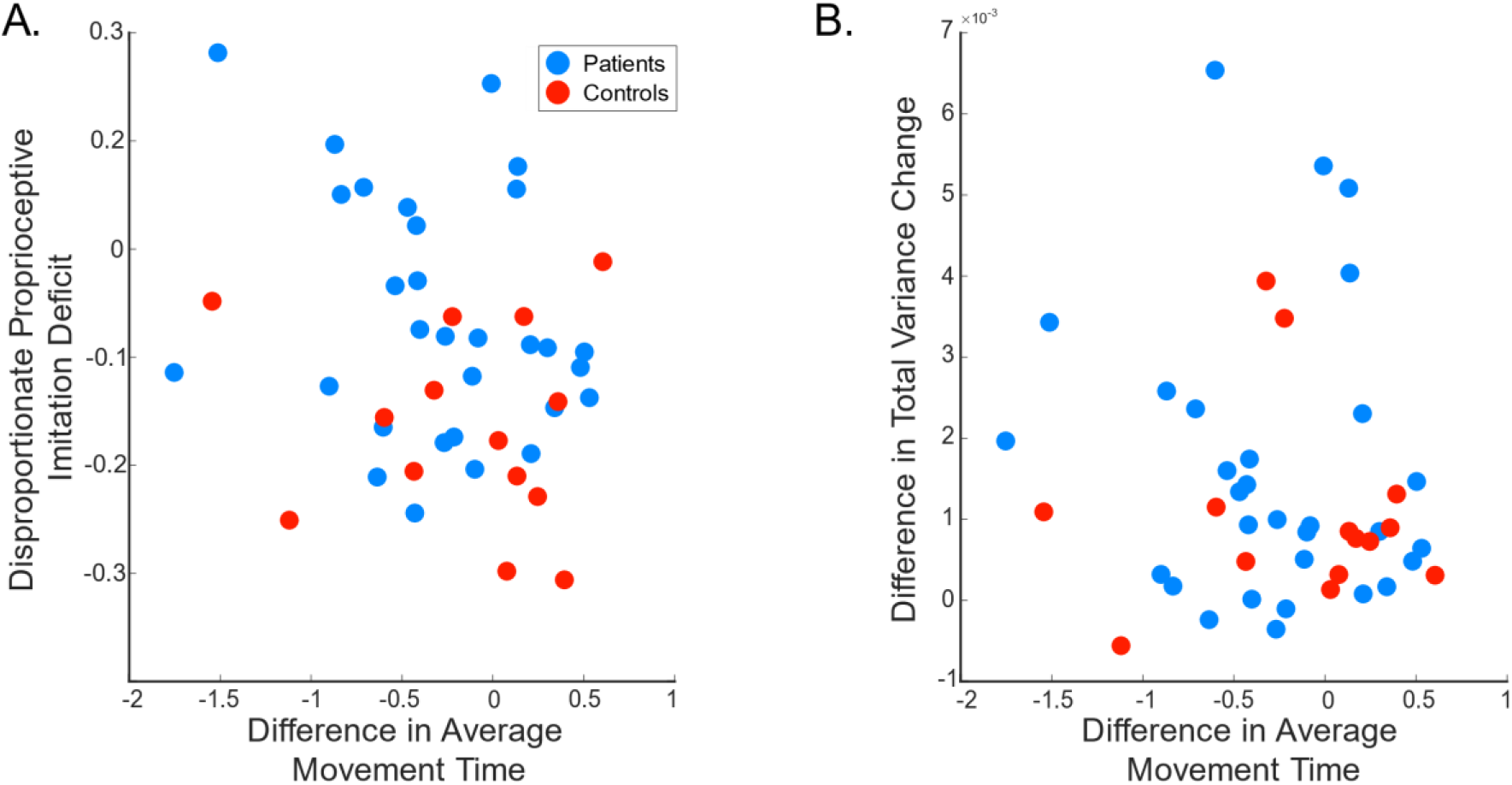
Imitation accuracy and consistency are not driven by a speed-accuracy trade-off. (A) Relationship of difference between average movement time from proprioceptive to visual blocks and disproportionate proprioceptive imitation deficits. No significant relationship was observed. (B) Relationship of difference between average movement time from proprioceptive to visual blocks and the difference between variability (change in Total Variation) between proprioceptive and visual imitation blocks. Again, no significant relationship was observed.

## 4 DISCUSSION

In this study we sought to explore the representations underlying the ability to imitate. In particular, we tested our Visual and Proprioceptive Hypothesis to see if we represent movement goals for imitation proprioceptively as well as visually; in other words, we assessed whether we specify both how our movements should feel as well as how they should look. We examined patients with left-hemisphere stroke, who are known to frequently exhibit imitation impairments associated with limb apraxia. We tested individuals under two conditions: one condition that provided only visual information about the cursor motion to be imitated and that encouraged reliance on visual feedback during imitation, and a second condition that provided only proprioceptive information both when cuing the desired movement and when imitating. We observed that in comparison to neurotypical controls, patients with left hemisphere stroke were indeed particularly impaired when asked to imitate movements that are cued purely proprioceptively. These imitation deficits were over and above what one would expect given these patients’ impairments at imitating movements cued visually (based on the performance of age-matched neurotypical controls). Importantly, we showed that these impairments could not be explained by underlying primary proprioceptive deficits, suggesting a higher-level problem with comparing incoming proprioceptive information to the movement goal. Moreover, our data are consistent with prior evidence that patients with apraxia are particularly reliant on visual rather than proprioceptive feedback when imitating (Howard et al., 2019; Jax et al., 2006): although visual and proprioceptive feedback were both available during the visual imitation condition of our task, we observed a strong relationship between performance in that condition and the visual trajectory-matching control task but not the proprioceptive control task. Together, our findings suggest that the ability to accurately imitate relies in part on the ability to represent how our desired movements ought to feel.

To our knowledge, while many have speculated on the importance of vision and proprioception for imitation, we are not aware of previous attempts to directly examine the separate roles of these two sensory modalities. Here, we have relied on two specific manipulations to better isolate these processes. First, analogous to our previous work (Wong et al., 2019), in the visual imitation condition we provided cues and feedback in the form of a cursor representing the hand position. This prior work demonstrated that cuing imitation in this way can expose imitation deficits akin to those exhibited when cuing imitation by watching an actor. By focusing the task solely on copying the motion of a cursor, and providing continuous cursor feedback of only that task-relevant dimension (i.e., end-effector position of the hand in the form of a cursor), we attempted to down-weight the contribution of proprioceptive guidance in this condition as no visual information about the positioning of the arm joints was available (Botvinick & Cohen, 1998; Graziano, 1999; Tanaka et al., 2009; Touzalin-Chretien et al., 2010). Individuals were generally able to perform this task, although as expected, patients did so with a range of performance accuracy.

The second important manipulation was the exclusion of all visual information in the proprioceptively cued trials. That is, we avoided any need to translate between or integrate vision and proprioception (a transformation that is necessary under the Kinesthetic-Visual Mapping hypothesis; Fig. 1) by letting participants feel the exact proprioceptive sensations they needed to reproduce in order to complete the imitation successfully. While it is certainly possible that individuals may still have transformed proprioceptive input into a visual movement percept (Mitchell, 2002), doing so would likely disrupt rather than aid performance (even assuming that the ability to perform these transformations was unimpaired) as it would introduce unrequired reference frame transformations, potential sensory integration biases, and possible spatial distortions that would be specific to the proprioceptive imitation condition (e.g. Burns & Blohm, 2010). Moreover, others have shown that reach-direction biases lie in different (unrelated) directions when reaching for targets that are specified visually or proprioceptively, suggesting that it is possible to represent a proprioceptive movement goal without necessarily representing it visually (Block & Bastian, 2010; Smeets et al., 2006). The fact that neurotypicals performed both visually-cued and proprioceptively-cued imitation with nearly equal accuracy (and that some neurotypicals even exhibited greater accuracy in the proprioceptive imitation condition) suggests these potential sources of errors in the proprioceptive condition are unlikely to have been introduced, and that the proprioceptive imitation condition indeed probed the ability to match proprioceptive feedback during an action to a proprioceptively-specified goal. Thus, any deficits observed in this condition were likely restricted to either the ability to properly locate the arm in space, or to represent the intended proprioceptive goal and compare it to incoming proprioceptive feedback. Our measure of proprioceptive accuracy allowed us to distinguish between these possibilities, and suggested the latter is more likely the case.

By comparing patients with apraxia against neurotypical controls in these conditions, we shed light on the direct contributions of proprioception in imitation and provide a more complete explanation of the way apraxia disrupts imitation ability. In particular, we found that patients were disproportionately worse at proprioceptive imitation compared to their neurotypical counterparts. When considered in conjunction with prior work showing that patients who are blindfolded exhibit greater imitation errors as well as fewer and less successful error-correction attempts (Howard et al., 2019; Jax et al., 2006), this suggests that proprioceptive abilities contribute to the imitation deficits observed in apraxia. Importantly, in our study we found no relationship between disproportionately poor proprioceptive performance and overall proprioceptive acuity deficits. While it is possible that patients were impaired at transforming between proprioceptive and visual representations, as we noted above such transformations may not necessarily be occurring in our task. Thus, the problem likely does not lie in individuals being unable to sense where their hand is when visual feedback is taken away. It is also important to recall that these assessments were performed on the ipsilesional side; thus, any observed performance deficits were unrelated to gross sensorimotor deficits (e.g., hemiparesis). Finally, we saw no evidence of an impairment related to gesture recognition. Instead, apraxia as measured by the imitation of meaningless visual gestures was associated with disproportionate difficulty with using proprioceptive information to guide imitation. We propose that gesture production deficits in limb apraxia are attributable in part to impairments in the ability to represent or access imitation goals specified in a proprioceptive format (i.e., how desired movements should feel). This deficit, in turn, leads to an overreliance on vision during imitation and poor performance imitating when blindfolded. Whether proprioceptive goals guide movements other than during imitation has not been tested, although there is some suggestion that this may indeed be the case (Kumar et al., 2021; Wong et al., 2012).

It is important to note that movements to be imitated are rarely cued proprioceptively; aside from hand-over-hand training (e.g., in sports or physical therapy), naturalistic imitation almost always involves viewing an action performed by someone else. We hypothesize that when watching others move, we are able to generate an estimate of the desired proprioceptive goal because we can observe how the body ought to be configured. However, there may also be other equally valid mechanisms for imitation that do not involve copying body positioning. For example, when imitating another species (e.g., dolphins using their whole body to copy a trainer’s hand), the only possible way to imitate is by imitating visual motion features such as spatial trajectories (Bauer & Johnson, 1994; Tayler & Saayman, 1973). We have previously demonstrated that even when cued visually by observing an actor, one can indeed imitate in two distinct ways: by attending to the body configurations of the actor or to the trajectory of the end-effector (Barhorst-Cates et al., 2020). Interestingly, patients with apraxia appear to be similarly impaired both when imitating actors or cursor trajectories, and this impairment correlates with a measure of meaningless imitation (Wong et al., 2019). It is surprising that we did not observe a similar correlation here, although this lack of correlation could reflect other less interesting task-related differences such as needing to control a robot in the horizontal plane. Nevertheless, in our current task, impairments in our visual-imitation condition (copying cursor motions that provided no body-configuration information) may have reflected difficulty in imitating end-effector trajectories, whereas impairments in our proprioceptive imitation condition (providing information about body motion) may have stressed body-configural imitation. The two conditions in our study – in addition to involving different sensory feedback and potentially different reference-frame transformations – may therefore have additionally evoked distinct mechanisms of imitation. It remains to be tested whether other commonly used measures of apraxia are related to the ability to copy trajectories or body positions.

Imitation impairments are a hallmark of apraxia, and are frequently associated with left-hemisphere strokes (Liepmann, 1905; Goldenberg, 2009, 2014; Goldenberg & Hagmann, 1997; Rothi et al., 1991; Tarhan et al., 2015). Indeed, previous work has suggested that imitation is subserved by specific loci in the brain, most often regions in the posterior parietal cortex (Buxbaum et al 2014; Caspers et al 2010). Interestingly, these regions have also been implicated in processing body-configural information (Chaminade et al., 2005; Graziano et al., 2000; Schwoebel et al., 2004) in a multimodal representation that includes proprioceptive information (Bolognini & Maravita, 2007; Làdavas, 2002; Limanowski & Blankenburg, 2017; Mountcastle et al., 1975), and are closely associated with nearby somatosensory cortex (Pandya & Seltzer, 1982; Whitlock, 2017). The posterior parietal cortex is also thought to play an important role in state estimation (i.e., a representation of how the body is currently positioned in space), which involves integration of visual and proprioceptive feedback as well as forward-model estimates (for review, see Medendorp & Heed, 2019; Shadmehr & Krakauer, 2008). Discrepancies between the current body state and the desired goal state are thought to be critical for driving motor corrections as well as learning (Diedrichsen et al., 2010; Zimmermann et al., 2012). Additionally, the supramarginal gyrus and arcuate fasciculus have been linked to both proprioceptive deficits (Findlater et al., 2018; Kenzie et al., 2016) and apraxia (Heilman et al., 1982). This overlap between brain regions implicated in imitation and proprioception suggests the presence of a network that is well positioned to represent proprioceptive goals for imitation.

Finally, although we did not observe a direct relationship between imitation ability and proprioceptive acuity, our findings hint that some patients with left-hemisphere strokes may exhibit primary proprioceptive deficits in the ipsilesional (“less affected”) arm, as seen by a few outliers in our proprioceptive control task. This complements the work of others that has argued for the presence of subtle ipsilesional deficits in stroke patients, including decreased dexterity and irregular reaches (Haaland et al., 1999; Schaefer et al., 2007, 2009; Sunderland et al., 1999). Such findings are important as attempts to assess various motor functions in stroke patients have often assumed that the ipsilesional side is intact. For example, some proprioception tasks entail passive movement of the ipsilesional arm and the instruction to match that movement with the contralesional arm (Dukelow et al., 2012; Semrau J. A. et al., 2015). The assumption is that any observed impairments solely reflect contralesional deficits. However, patients’ performance on our proprioceptive control task suggests that this need not be true, and supports the importance of directly assessing sensory deficits even on the ipsilesional side. In addition, it is possible that patients also exhibit subtle deficits in transforming between visual and proprioceptive representations, or in transforming between allocentric and egocentric reference frames – factors that could additionally contribute to the imitation impairments we observed here. Future work is needed to understand if such impairments exist, how they arise, and whether these effects are also left-lateralized along with apraxia.

In summary, here we have presented evidence that an important step in motor planning is to represent how our body should feel when performing that movement. When it comes to imitation, this means transforming observed movements performed by others into an estimate of how those movements should feel when performed by ourselves; this allows us to then directly compare proprioceptive feedback of our limb to a goal describing what that feedback should be, and thereby detect and correct errors in our movement. Under typical circumstances when vision is also available, it is likely that this proprioceptive goal is used in combination with or superseded by a representation of how our movement should look visually. What remains unclear is whether we combine visual and proprioceptive information to form a single (multimodal or amodal) representation of our movement goal to which we can compare a state estimate of our current body position based on visual, proprioceptive, and feedforward predictions (and if so, to what degree vision and proprioception are weighted to form such a representation), or if these exist as separate unimodal representations. Nevertheless, our findings suggest an important contribution of proprioceptive goals to the ability to imitate meaningless movements, and that disruption of this ability contributes to imitation impairments in patients with apraxia following left-hemisphere stroke. It follows that patients with such deficits may attempt to rely on visual feedback in a compensatory manner. Of future interest is the question of whether these patients could be instead taught to adopt an entirely different imitation strategy, such as focusing on planning end-effector trajectories rather than body configurations.

## ACKNOWLEDGEMENTS

This work was supported by grant AES 19-02 from the Albert Einstein Society of the Einstein Healthcare Network, NIH grant R01 NS115862 to ALW, and NIH grant R01 NS099061 to LJB.

## Footnotes

1 We observed one outlier patient whose imitation accuracy in the easy proprioceptive-imitation condition was much worse than any other individual in either the easy or hard difficulty conditions (including their own performance in the hard condition; Fig. 3A). To check if this participant was skewing our findings, we re-ran all of our models (including the models in the sections below) without this participant but saw no qualitative change in our findings (statistics not reported); we therefore retained this participant in our dataset.

